# Polyphosphate affects cytoplasmic and chromosomal dynamics in nitrogen-starved *Pseudomonas aeruginosa*

**DOI:** 10.1101/2021.12.23.473106

**Authors:** S. Magkiriadou, A. Habel, W. L. Stepp, D. K. Newman, S. Manley, L. R. Racki

## Abstract

Polyphosphate (polyP) synthesis is a ubiquitous stress and starvation response in bacteria. In diverse species, mutants unable to make polyP have a wide variety of physiological defects, but the mechanisms by which this simple polyanion exerts its effects remain unclear. One possibility is that polyP’s many functions stem from global effects on the biophysical properties of the cell. We characterize the effect of polyphosphate on cytoplasmic mobility under nitrogen-starvation conditions in the opportunistic pathogen *Pseudomonas aeruginosa*. Using fluorescence microscopy and particle tracking, we characterize the motion of chromosomal loci and free tracer particles in the cytoplasm. In the absence of polyP and upon starvation, we observe an increase in mobility both for chromosomal loci and for tracer particles. Tracer particles reveal that polyP also modulates the partitioning between a ‘more mobile’ and a ‘less mobile’ population: small particles in cells unable to make polyP are more likely to be ‘mobile’ and explore more of the cytoplasm, particularly during starvation. We speculate that this larger freedom of motion may be a consequence of nucleoid decompaction, which we also observe in starved cells deficient in polyP. Our observations suggest that polyP limits cytoplasmic mobility and accessibility during nitrogen starvation, which may help to explain the pleiotropic phenotypes observed in the absence of polyP.

## Introduction

While most of our understanding of bacterial physiology comes from studying rapidly growing cells, growth arrested states have deep relevance for natural and clinical settings. Bacterial growth rates can be extremely heterogeneous in the context of chronic bacterial infections, and slow- or nongrowing cells may have an increased likelihood of tolerance to conventional antibiotics. Distinct molecular activities underlie fitness and survival during growth arrest, as bacteria undergo metabolic, regulatory, and biophysical changes in response to nutrient limitation (1). For example, the nucleoid experiences a decrease in supercoiling and an increase in chromosome compaction(2–4). These changes are thought to be driven by decreased transcription and increased synthesis of starvation-specific nucleoid associated structural proteins (NAPs) (5–8). Meanwhile, the cytoplasm exhibits an overall decrease in volume and increase in density in some bacteria as well as in yeast (9, 10). In some cases, changes in the dynamics of cytoplasmic components have also been observed (10, 11). For example, during carbon starvation, *Caulobacter crescentus* exhibits decreased or arrested cytoplasmic transport, especially pronounced for larger complexes, leading to a proposed model whereby metabolic activity is responsible for fluidizing the cytoplasm (11).

Another dramatic change to cellular organization that occurs in many bacteria during starvation is the production of polyphosphate (polyP). Bacteria spend ATP to synthesize polyP polymers consisting of long chains of phosphoryl groups, which can come together to form granule superstructures. Making polyP is important for starvation survival, with defects in polyP synthesis leading to pleiotropic fitness effects (12, 13). In *Pseudomonas aeruginosa*, PolyP granules form in the nucleoid region of the cell during the first few hours of nutrient starvation and persist into deeper starvation. It also appears that making polyP can functionally affect the chromosome: polyP modulates cell cycle progression in *E. coli* and *C. crescentus*, and in *P. aeruginosa* it is required for efficient cell cycle exit in response to nitrogen starvation (14–16). Furthermore, *P. aeruginosa* cells that cannot make polyP activate the SOS DNA damage response, suggesting that polyP may have a protective function during starvation (16). However, the impact of polyP on the biophysical properties of cells, including chromosome organization and cytoplasmic dynamics, remains unknown.

In this study, we characterize the effects of polyP on the biophysical properties of the chromosome and the cytoplasm during the crucial transition period between exponential growth and deep starvation, the period during which polyP production ramps up, as evidenced by the formation of polyP granules. We focus on nitrogen starvation, a condition in which we have previously characterized polyP granule biogenesis (16). At 6 hours of nitrogen limitation, we find that the mobility of the chromosome and large cytoplasmic complexes in WT cells are similar to those we observe under exponential growth conditions. In contrast, cells unable to make polyP exhibit a significant increase in chromosomal and cytoplasmic mobility, concomitant with decompaction of the nucleoid. The dramatic polyP-dependent effects we observe on cytoplasmic transport properties occur under nitrogen starvation, but not carbon starvation, suggesting that polyP may have distinct functions under different types of starvation.

## Results

### Chromosome mobility under nitrogen starvation

We measured chromosome motion in strains engineered to contain the ParS^pMT1^ DNA binding sequence at a locus 19.5kb from the chromosomal origin of replication, and expressing its cognate DNA binding protein GFP-ParB^pMT1^. When bound by ParB^pMT1^, which forms an oligomer, the ParS^pMT1^ site appears as either one or two bright punctae, depending on the stage of chromosomal replication (Fig. 1A). We recorded the motion of these fluorescently labeled loci with widefield microscopy at 0.2 frames/s and extracted their subpixel locations in each frame using a Gaussian fitting algorithm (17, 18). We then connected the positions of each locus between frames based on proximity, to form trajectories. For each trajectory we characterized the dynamics of the chromosomal locus through its mean squared displacement, defined as:

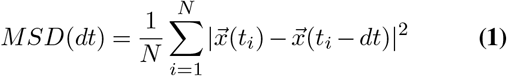

**Fig. 1.**
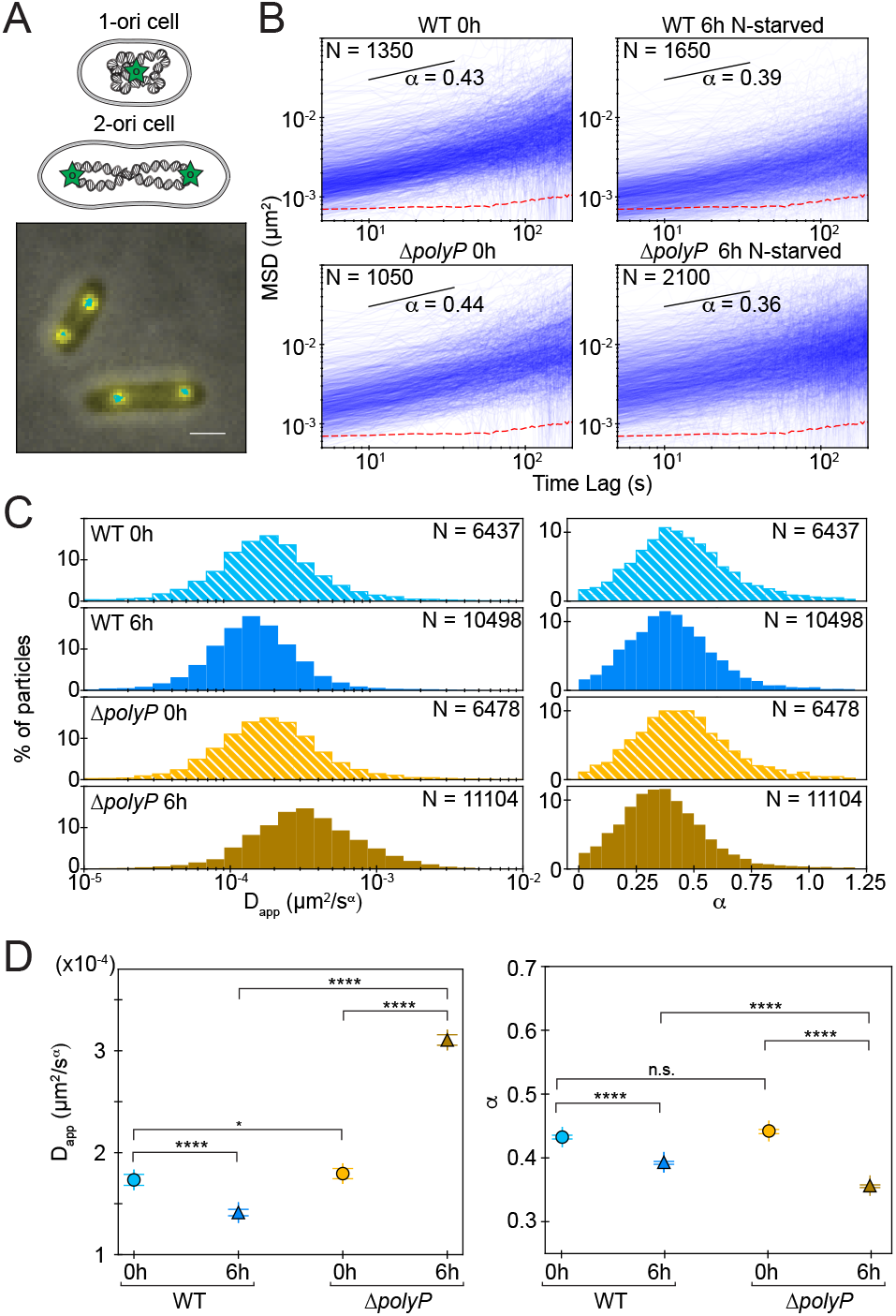
Polyphosphate modulates the mobility of chromosomal origins during nitrogen starvation. (A) Top: We label the origins of the bacterial chromosome with GFP (green stars) as described in the Supplementary Information. Depending on their stage in the cell cycle, some cells have one origin while others have two. Bottom: two example cells with their chromosomal origins labeled (yellow spots) and tracked in time (cyan lines). Scale bar: 1 μm. (B) Log-log plots of example mean-squared-displacement curves for a few hundred chromosomal origins at four conditions: WT cells (top row) and ΔpolyP cells (bottom row), 0 h (left column) and 6 h into N-starvation (right column). The *α* value shown in each panel is the median of each distribution. The black line in each panel represents the slope that corresponds to the quoted *α* value, obtained together with *D_app_* from linear fits over the time interval spanned by the black line (10s – 40s). N is the number of plotted curves; we show a randomly chosen subset for clarity. (C) Histograms of *D_app_* (left column) and *α* (right column) for chromosomal origins in WT (blue) and ΔpolyP (orange) cells, 0 h (light colors, striped) and 6 h into N-starvation (dark colors, solid). N is the total number of spots at each condition. (D) Medians of the *D_app_* (left) and *α* (right) distributions for each condition. The error bars denote the standard error of the mean, and the asterisks denote the statistical significance of the differences in the medians according to Mood’s median test (see SI).

where 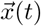 is the locus position at time *t*, *dt* is the time lag between measured positions, and *N* is the number of statistically independent measurements taken along the trajectory for that locus and time lag (17).

To investigate the effects of starvation and polyphosphate, we quantified chromosomal dynamics in both wild-type cells (WT) and mutant cells lacking the enzymes required to make polyP (ΔpolyP). We performed our measurements under two conditions: under exponential growth in MOPS-buffered minimal media (0 h), or under starvation, 6 hours after transferring cells to nitrogen-limited minimal media (6 h). We note that while the agarose pads we used for imaging contained no added nitrogen, we observed a small amount of growth over time on the pads. We characterized the effect of the amount of time cells sat on agarose pads before imaging to confirm that this growth did not affect our measurements (Fig. S1). For all conditions, *MSD* curves appear linear on a log-log scale, consistent with a power-law scaling (Fig. 1B). Whereas purely diffusive motion is described as a random walk, captured by the Smoluchowski-Einstein relation (19) and a slope of 1, we observe lower slopes typical of sub-diffusive motion.

To further quantify and interpret this motion, we consider the general model:

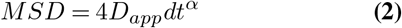

Here the scaling exponent, *α*, corresponds to the slope on a log-log plot, while the apparent diffusion coefficient, *D_app_*, is 1/4 the value of the *MSD* at *dt* = 1 s. From each individual chromosomal locus’ *MSD* curve, we extracted *D_app_* and *α* by fitting over the time interval 10-40 s. This ensured that the curves lie above the measurement noise and *N* is large enough to offer low statistical noise (see Methods).

We measured the chromosomal motion of many individual cells, and find that the resulting histograms of *D_app_* and *α* for all four conditions are well-described by a single-mode distribution (Fig. 1C). Before the onset of starvation (0 h), distributions in both WT and ΔpolyP cells have similar median values and widths (Fig. 1C, D). In both, we find a distribution of *D_app_* centered at 1.7 − 1.8 × 10^*−*4^*μm*^2^/*s^α^* and a distribution of *α* centered at 0.43 − 0.44. Compared with previous reports on chromosomal loci in *E. coli*, these median values of *D_app_* are about 10-fold lower (20) while *α* values are consistent (21). In contrast, after nitrogen starvation (6 h), while in WT cells *D_app_* decreases slightly to 1.4 × 10^*−*4^*μm*^2^/*s^α^*, in ΔpolyP cells *D_app_* approximately doubles at 3.1 × 10^*−*4^*μm*^2^/*s^α^*, indicating that chromosomal loci experience higher displacements. At the same time, although the distributions of *α* for both strains show significant overlap, the median value of *α* decreases slightly to 0.39 for WT cells and to 0.36 for ΔpolyP cells.

### Cytoplasmic mobility under nitrogen starvation

We wondered whether the differences we observed in the motion of chromosomal origins are limited to the nucleoid, or whether they could be reflective of more wide-ranging changes to the cytoplasm. To investigate cytoplasmic mobility, we adapted GFP-μNS probe particles previously developed for *E. coli* and *C. crescentus*, for *P. aeruginosa*. We induced expression of GFP-fused μNS, an avian reovirus protein expected to have no specific interactions with components of the bacterial cytoplasm (11), in both WT and ΔpolyP strains (see Methods). We observed fluorescent punctae, indicating that GFP-μNS proteins were assembling into particles.

We observed that GFP-μNS particles are more mobile than chromosomal origins; thus, to accurately capture their motion we recorded time-lapse movies at a higher frame rate (4.76 frames/s). As with chromosomal loci, we localized and then tracked the GFP-μNS particles (17, 18). Qualitatively, we noticed two types of motion: some particles appeared locally confined, while others explored the cytoplasmic volume more freely (Fig. 2A). We calculated the *MSD* following Equation [1] as before. *MSD* curves appear to follow a powerlaw dependence on time lag as found for chromosomal origins; however, particles take on a wider range of values and slopes, consistent with our qualitative observation (Fig. 2B). Furthermore, in the case of nitrogen-starved ΔpolyP cells, we clearly discern two modes of motion: a lower mobility mode in which particles display an *MSD* near the measurement noise even at the longest time lags, and a higher mobility mode in which particles are able to move further through the cell as time passes.

**Fig. 2.**
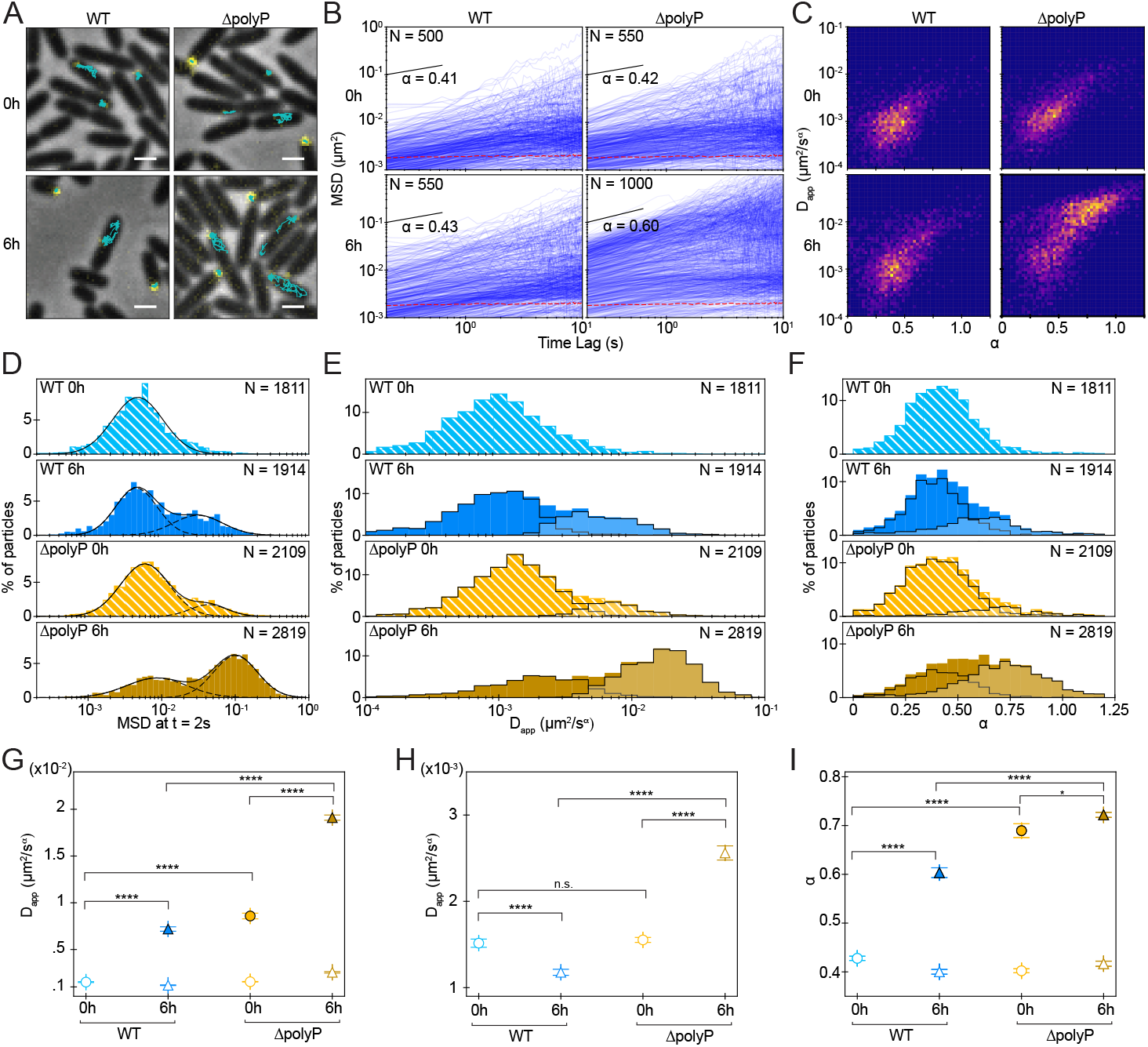
Polyphosphate affects cytoplasmic mobility during nitrogen starvation. (A) Examples of cells with fluorescently labeled μNS particles (yellow spots) and their tracks in time (cyan lines). In both the WT (left column) and the ΔpolyP strains (right column) we observe two types of behavior at 0 h (top row) and 6 h into N-starvation (bottom row): some μNS particles appear confined and others explore a larger area of the cytoplasm. Scale bar: 1 μm. (B) Examples of mean-squared-displacement curves for a randomly chosen subset of particles in each condition. The *α* values in each panel denote the median of the distribution; they are obtained from linear fits over the time interval spanned by the black line (0.21 s – 0.84 s). N is the number of plotted curves; we show a randomly chosen subset for clarity. The presence of two types of behavior is best visible for ΔpolyP cells 6 h into N-starvation (bottom right panel), where the *MSD* curves split into two branches. The red dotted line indicates our empirical noise floor, measured as the ensemble mean-squared-displacement of μNS particles in fixed cells. (C) Heatmaps of *D_app_* vs *α* for WT (left column) and ΔpolyP cells (right column) at 0 h (top row) and 6 h into N-starvation (bottom row). (D) Histograms of the mean-squared-displacement values at t = 2.1 seconds. Except for WT at 0 h, which fits well to a normal distribution (black continuous line), all other conditions are well described by a binormal fit (black continuous lines). The two modes that comprise the bimodal distributions (black dashed lines) correspond to the two branches of lines in (B). (E) Histograms of *D_app_* values extracted from the *MSDs* shown in (B). The black outlines mark the two sub-distributions that comprise the full distribution, one of particles of low mobility and one of particles of high mobility (lighter color); here the type of mobility is defined by the value of the particle’s *MSD* at 2.1s, shown in (D). (F) Histograms of *α* values extracted from the *MSDs* shown in (B), with the two sub-populations of mobility shown as in (E). In (D),(E),(F), N indicates the total number of particles. (G),(H),(I) Mean values for *D_app_* (G,H) and *α* (I) for each mobility mode in each condition. The error bars denote the standard error of the mean. Open shapes denote ‘low-mobility’ mode, and filled shapes denote ‘high-mobility’ mode. The asterisks denote the statistical significance of the differences in the means according to the T-test (see SI).

To quantify these sub-populations of GFP-μNS particles and link them with the qualitative observation of different degrees of confinement, we extracted the values of the *MSD* curves at a time lag *dt* = 2.1*s*, *MSD*(*dt* = 2.1*s*) (Fig. 2D). The histogram of *MSD*(*dt* = 2.1*s*) for the unstarved (0 h) WT strain shows a single mode. In contrast, for the nitrogen-starved WT and ΔpolyP mutant strains, as well as for the unstarved (0 h) ΔpolyP mutant, we observe two particle sub-populations that are well fit as bimodal distributions. Taking as a boundary the point of intersection between the two curves that we obtain from the bimodal fits, we estimated the percentage of low- and high-mobility particles under each condition. We find that for WT at 6 h and for ΔpolyP at 0 h a minority of GFP-μNS particles occupy the high-mobility mode (31% and 15% respectively), while for the ΔpolyP mutant at 6 h more particles occupy the high-mobility mode (62%), as summarized in Table 1. In this context, the single mode of the WT strain at 0 h corresponds to low-mobility particles.

**Table 1.** Distribution of μNS particles into modes of low and high mobility.

| condition | % in low-mobility mode | % in high-mobility mode |
| --- | --- | --- |
| WT 0 h | 100 | 0 |
| WT 6 h | 69 | 31 |
| ΔpolyP 0 h | 85 | 15 |
| ΔpolyP 6 h | 38 | 62 |

We further quantified the motion of μNS particles via the apparent diffusion coefficient, *D_app_*, and the scaling exponent, *α*, which we obtained from fits of their *MSD* over the time interval 0.21 < *dt* < 0.84 sec. We find that *α* correlates positively with *D_app_* (Fig. 2C), indicating that particles whose motion is less subdiffusive also explore more of the cell. We assigned each particle to a mobility group defined by its *MSD*(*dt* = 2.1*s*) belonging in either the low mobility or high mobility mode (Fig. 2D), and examined the distributions of *D_app_* and *α* (Fig. 2E,F). Although the modes are less distinctly separated than in the *MSD*(*dt* = 2.1*s*) histograms, they remain as well-defined sub-populations. For each mode we define a mean value under each condition (Fig. 2G,H,I). Interestingly, the two modes are differentially affected by the different conditions. For the low mobility mode, the mean value of *D_app_* is similar for WT at 0 h (1.5 × 10^*−*3^*μm*^2^/*s^α^*) and at 6 h (1.2 × 10^*−*3^*μm*^2^/*s^α^*) and for ΔpolyP at 0 h (1.6 × 10^*−*3^*μm*^2^/*s^α^*), while in the nitrogen-starved ΔpolyP mutant, *D_app_* is higher by about a factor of 2 (2.6 × 10^*−*3^*μm*^2^/*s^α^*). This quantitatively repeats the trends observed for *D_app_* of the chromosome. The scaling exponents of the low mobility modes remain at similar values in all conditions (*α* ~ 0.4) and are also similar to those observed for the chromosome.

By contrast, for the high mobility mode, *α* is closer to the free diffusion value of 1, with a value of 0.6 for the WT at 6 h, 0.69 for the ΔpolyP mutant at 0 h, and 0.72 for the ΔpolyP mutant at 6 h. For this mode, WT at 6 h and ΔpolyP at 0 h are again comparable in *D_app_* (7.2 × 10^*−*3^*μm*^2^/*s^α^* for WT 6 h and 8.6 × 10^*−*3^*μm*^2^/*s^α^* for ΔpolyP at 0 h). In comparison, the mutant is clearly distinct under starvation, with a mean *D_app_* that is 3-fold higher (19 × 10^*−*3^*μm*^2^/*s^α^*).

### Cytoplasmic mobility under nitrogen starvation and particle size

As GFP-μNS particles move, they probe the cytoplasm, offering insights into its transport properties. Moreover, GFP-μNS particles come in a variety of sizes, since monomers can assemble in different stoichiometries (11); this allows us to probe the cytoplasm overa a range of scales. We estimated the relative sizes of individual particles from the integrated intensity of their fluorescent signal (see Methods). In a viscous medium, the diffusion coefficient of spherical particles scales inversely with particle size as described by the Stokes-Einstein equation:

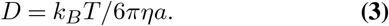

Here the numerator represents the thermal energy, which is the product of the Boltzmann constant *k_B_* and the temperature *T*, and the denominator represents the viscous dissipation, as the product of the viscosity of the medium *η*, the particle radius *a*, and a geometric constant. Considering that the bacterial cytoplasm is a complex, viscoelastic, and heterogeneous medium, we do not expect this precise scaling to apply to our GFP-μNS particles. Still, departures from this scaling, and the dynamic behavior of particles of different sizes may provide insights into the different mobility modes we observe in the overall population of particles (Fig. 2).

Cells in all strains and conditions we measured contained particles with a similar size distribution (Fig. S2 A); thus, their mobility distributions can be directly compared. As expected, particle size correlates inversely with *D_app_*; smaller particles generally diffuse faster than larger ones in all conditions (Fig. 3A). Particle size also has a weak effect on the subdiffusive scaling exponent (Fig. 3B); while we measured low values (*α* < 0.5) for a wide range of particle sizes, higher values (*α* > 0.5) closer to free diffusion are more often reached by smaller particles. To examine how size affects particle mobility in the cytoplasm, we divided particles into five groups according to their fluorescence intensity. Small particles fall into the dimmest group, “Bin 1,” medium particles into the middle group, “Bin 3,” and large particles into the brightest group, “Bin 5.” Examining the WT cells at 0 h, we observe that while *D_app_* decreases with size (Fig. 3A), *α* remains constant (Fig. 3B). For the remaining conditions, we maintained the labels on individual particles of “low mobility” or “high mobility” described above (Fig. 2). The low mobility population of particles in WT cells at 6 h nitrogen starvation and in ΔpolyP cells at 0 h have similar behavior: increasing size signifies a lower *D_app_* and unchanged *α*. In these two conditions the high mobility particles are found nearly exclusively among the small particles, and the values of *D_app_* appear less dependent on size. In the nitrogen-starved ΔpolyP mutant, both mobility populations are clearly visible not only for small particles, but also for medium and large ones. The dominance of the high mobility mode for the smallest particles is especially striking.

**Fig. 3.**
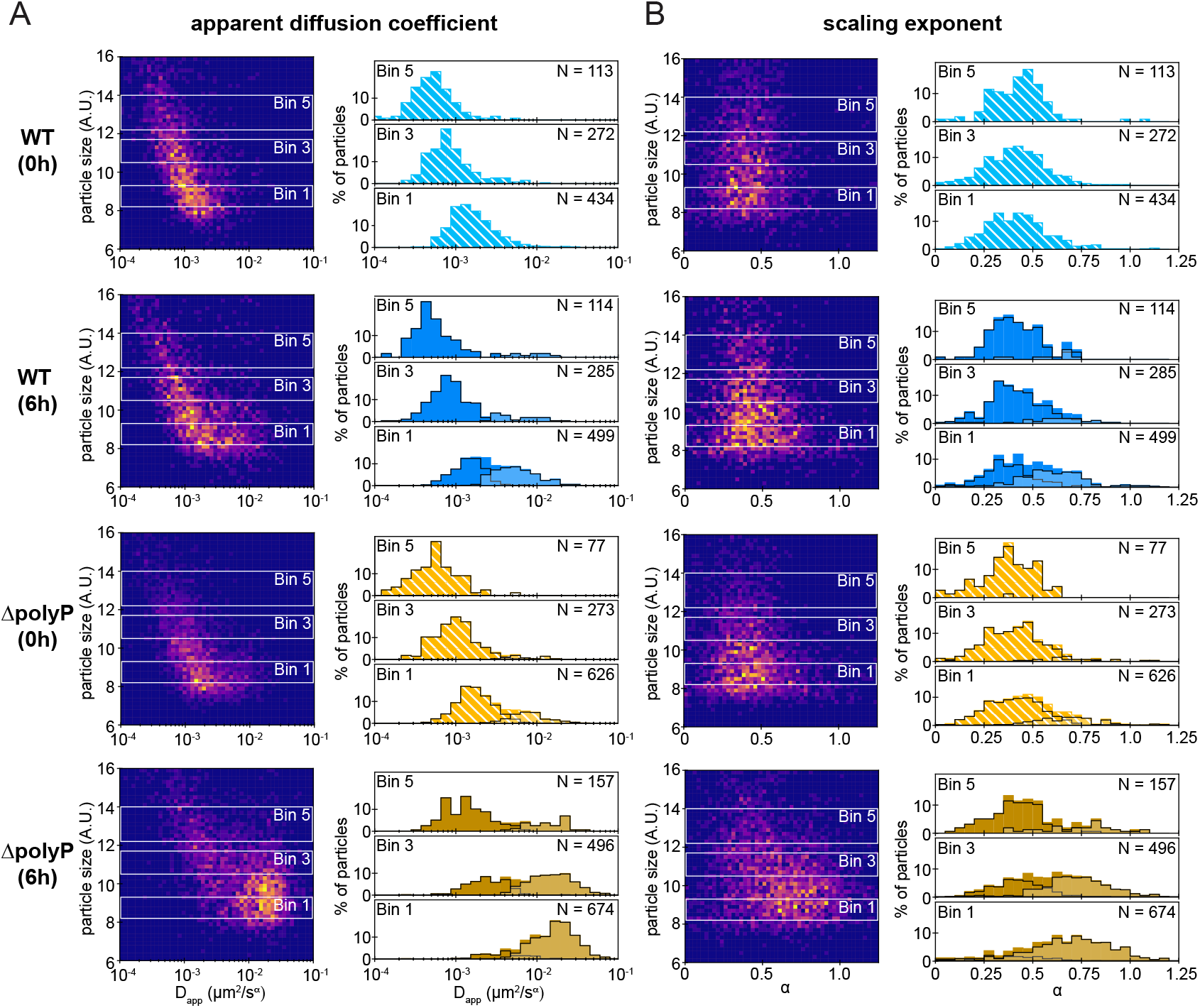
The effect of polyphosphate on cytoplasmic mobility 6 h into nitrogen starvation depends on particle size. (A) Left: heatmap of particle size vs *D_app_* ; right: histograms of the *D_app_* distribution for three size bins in each condition, colored as in Fig. 2(E). (B) Left: Heatmap of particle size vs *α*; right: histograms of the *α* distribution for three size bins in each condition, colored as in (A). In (A), (B), bin 1 corresponds to small particles, bin 3 to medium-sized particles, and bin 5 to large particles.

### Cytoplasmic mobility under carbon starvation

We wondered whether the increased mobility we observed in ΔpolyP cells was specific to nitrogen starvation or if it might also be observed under carbon starvation, a condition shown for some bacterial species to suppress cytoplasmic motion (11). Under carbon starvation we observe broadening of the distribution of *D_app_* values, with a significant fraction of particles becoming less diffusive than in unstarved cells, but also some particles exhibiting increased mobility compared to unstarved cells (Fig. S3 B). We also observe for all particles a shift to lower *α* values, including the emergence of a large subpopulation of cells with *α* values of <0.25, indicating increased confinement (Fig. S3 C). Binning of the particles by size reveals that, as with nitrogen starvation, smaller particle size correlates with increased mobility (Fig. S4) We observe a similar effect on *D_app_* and *α* in the ΔpolyP mutant during carbon starvation, without the emergence of a more mobile population in ΔpolyP compared with WT cells. Therefore, polyP does not appear to have a significant effect on cytoplasmic mobility during carbon starvation, a condition under which transport is thought to be affected by energy limitation.

### Nucleoid occupancy of the cytoplasm under nitrogen starvation

The cellular interior is a crowded and structured environment whose organization and occupancy can strongly affect motion through it. Cytoplasmic mobility has been shown in *E. coli* and *C. crescentus* to depend both on the percentage of cytoplasmic area occupied by the chromosome, or “nucleocytoplasmic ratio” (NC ratio) (22), and on the effective ‘mesh size’ imposed by the chromosomal DNA within the nucleoid region (23). To elucidate the role of the nucleoid in our measurements of cytoplasmic mobility, we labeled the chromosome by expressing a fluorescent chimera of the nucleoid associated protein HU, which globally binds the chromosome, in WT and in ΔpolyP mutant backgrounds. We expressed HU-mCherry from a second copy of its native promoter integrated at the *attB* site on the chromosome. We then performed nucleoid and cell segmentation and quantification (Methods, Fig. 4A, Supplementary Information) as previously described (22, 24).

**Fig. 4.**
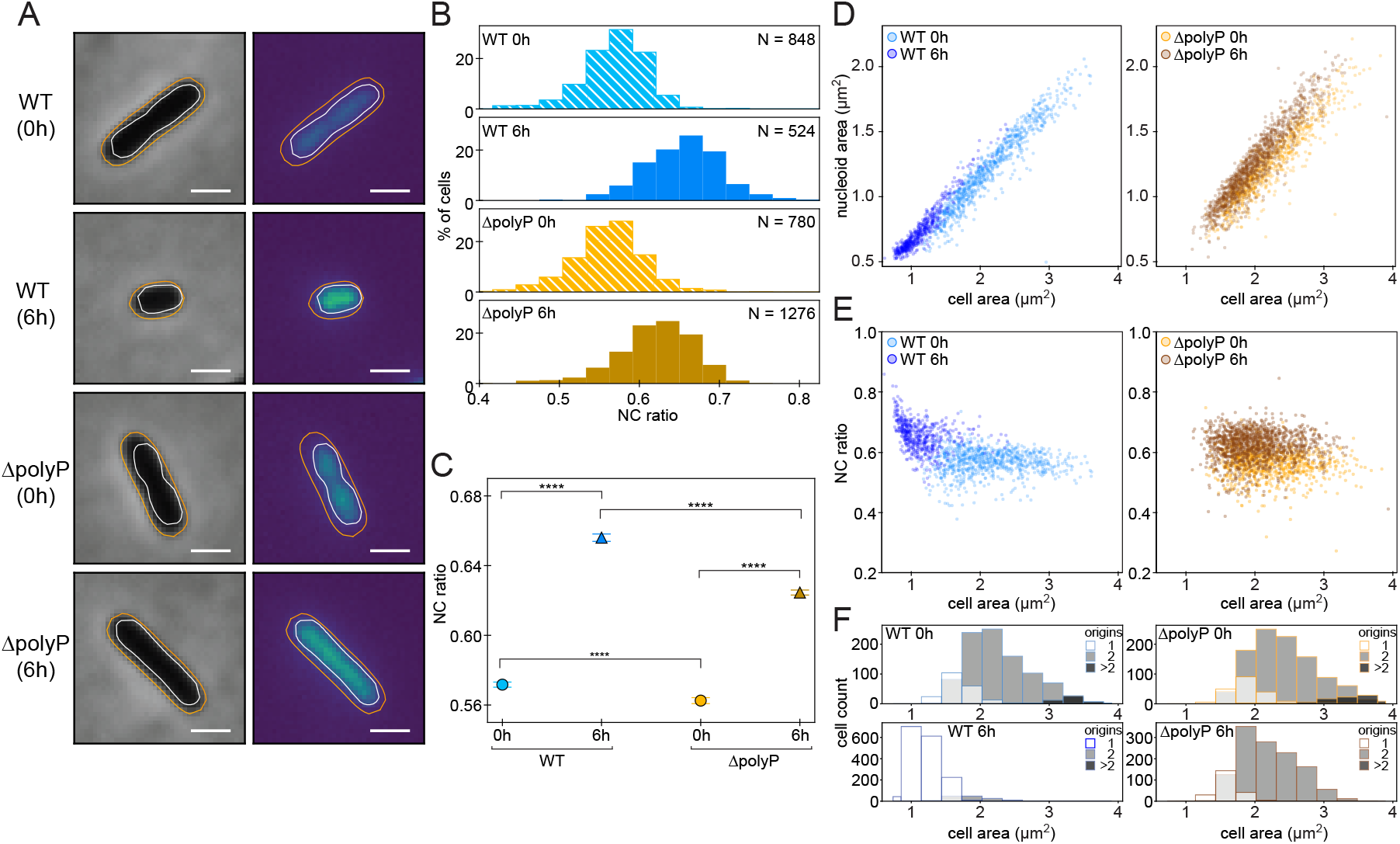
Polyphosphate affects nucleoid occupancy of the cytoplasm. (A) Examples of cells whose chromosome has been labeled with HU-mCherry. Left column: phase channel; right column: fluorescence channel. We segmented the cells (orange outlines) and the nucleoid (white outlines) to measure the ratio of the area occupied by the chromosome to the area of the whole cell (NC ratio). Scale bar: 1 μm. (B) Histograms of the NC ratio for all four conditions. (C) Medians of the distributions of the NC ratios shown in (B). The error bars denote the standard error of the mean and the asterisks denote the statistical significance of the differences in the medians according to the Mood’s median test (see SI). (D) Scatter plots of nucleoid area vs cell area. (E) Scatter plots of the NC ratio vs cell area. (F) Stacked histograms of the cell area, grouped by the number of chromosomal origins per cell.

Before the onset of starvation, the distributions of NC ratios for both WT and ΔpolyP strains are similar in form and in median values (~ 0.57 for the WT, ~ 0.56 for ΔpolyP, Fig. 4B,C). The total area occupied by the nucleoid is also similar (Fig. 4D). Six hours into nitrogen starvation the distribution of the NC ratios shifts to markedly higher values for both strains (median NC ratio ~ 0.66 for WT and ~ 0.63 for ΔpolyP cells, Fig. 4B,C,E). However, WT cells are significantly smaller during starvation than are ΔpolyP cells (Fig. 4A), and their total DNA content also differs. While most WT cells have finished the cell cycle and have a single chromosome per cell at 6 h, most ΔpolyP cells have not exited the cell cycle and a similar fraction of cells has two origins compared to both WT 0 h and ΔpolyP 0 h cells (Fig. 4F). The total cytoplasmic area taken up by the nucleoid for cells of all sizes is greater in ΔpolyP 6 h cells than for both WT and ΔpolyP growing cells at 0 h (Fig. 4D). We note that the total fluorescence intensity of the HU-mCherry chimera is brighter for both WT and ΔpolyP size-matched cells during nitrogen starvation (Fig. S5 A). We used Sybr Green labeling of the chromosome as an independent method to characterize nucleoid area and the NC ratio, for which we observe a similar increases in the ΔpolyP mutant as we found using HU-mCherry (Fig. S5 C,D,E,F). In summary, while cell size and the number of chromosomes per cell is similar in growing WT, growing ΔpolyP, and starved ΔpolyP cells, the DNA occupies more of the cytoplasm in starved ΔpolyP cells than in either growing WT cells or growing ΔpolyP cells. This suggests that the chromosome is relatively decompacted in ΔpolyP cells, and conversely, that polyP may play a role in chromosome compaction during nitrogen starvation.

## Discussion

### PolyP suppresses diffusion during nitrogen starvation

Previous work has shown that properties of the bacterial cytoplasm can change in response to energy depletion (11). Energy depletion offers a natural mechanism to decrease cytoplasmic motion under carbon starvation, but tuning cytoplasmic mobility under other forms of growth arrest may be important for regulating many processes. Here we discover that polyP affects the properties of the cytoplasm during nitrogen starvation. We observe an increase in the apparent diffusion coefficient for the chromosome as well as for free particles in ΔpolyP cells, indicating that polyP suppresses the motion of free particles as well as of chromosomal loci during nitrogen starvation. While the values of *D_app_* are different for the chromosome and for free particles as expected from their different physical properties, the scaling of this effect is similar, about 2-fold increase in the absence of polyP for both.

We speculate that polyP modulates diffusion by some combination of two biophysical mechanisms: a direct effect of the polyP polymer on the viscosity of the cytoplasm, and an indirect effect on cytoplasmic bioenergetics via nucleotide pools. Viscoadaptation during stress and starvation was recently shown to be driven by polymers and small molecules such as glycogen and trehalose in yeast (25). We speculate that the polyP polymer might serve a similar a role in bacteria. In addition to viscoadaptation, another non-exclusive mechanism may be a direct effect on cellular bioenergetics via ATP. PolyP, which is synthesized by polyP kinase enzymes from ATP, may decrease available ATP pools, as has been suggested in studies in both *E. coli* during oxidative stress and *Chlamydomonas reinhardtii* during nitrogen starvation (26, 27). Increased ATP would be expected to lead to more frequent cytoplasmic rearrangements and micro-agitations. ATP’s putative function as a hydrotrope, enhancing overall protein solubility, could also modulate overall cytoplasmic motion (28).

### PolyP constrains the apparent mesh size of the cytoplasm during nitrogen starvation

In addition to a global affect on *D_app_*, we find that polyP modulates the partitioning of GFP-μNS particles between ‘mobile’ and ‘immobile’ populations during nitrogen starvation. We observed in WT cells the emergence of a small population of particles during nitrogen starvation that are more mobile than during exponential growth. These particles have both a larger *D_app_* and a larger scaling factor, indicating that they experience less confinement. In the ΔpolyP mutant, the ‘mobile’ fraction predominates during nitrogen starvation. This partitioning is not an artifact of differences in particle sizes in WT vs ΔpolyP cells, since the size distributions in the two strains are similar. For the largest particle bin size both WT and ΔpolyP cells are mostly ‘immobile.’ For the intermediate particle bin size WT cells have almost no mobile particles, while ΔpolyP cells have equal fractions of mobile and immobile particles; thus at this length scale the absence of polyP results in more accessibility. Finally, for the smallest particle bin size both strains have mobile particles, but ΔpolyP cells contain a nearly entirely high mobility population, whose diffusivities are even higher than those of the mobile particles in WT cells. These observations reveal that intermediate and small particles are freer to explore the cytoplasm in ΔpolyP cells, particularly during starvation. In turn, this suggests that the structuring of the cytoplasm during nitrogen starvation depends on polyP. A possible explanation for the partitioning we observe would be that cells unable to make polyP exhibit a larger effective ‘mesh size’ than WT cells.

To further explore possible mechanisms that might underpin the effects of polyP on chromosomal and cytoplasmic motion, we turned to the nucleoid. The nucleoid is thought to be a major contributor to the effective mesh size of the cytoplasm, and it has been shown in some bacterial species and conditions to compact during starvation (4). Cell volume in WT *P. aeruginosa* cells also decreases, which may contribute to overall crowding, and increased cytoplasmic density has been reported in *E. coli* under some starvation conditions (9). We have previously shown that ΔpolyP cells fail to exit the cell cycle during nitrogen starvation, and therefore remain as large as WT growing cells (16). If the chromosome did not change in its compaction, we would expect therefore to see a similar mesh size for ΔpolyP cells as WT growing cells. To our surprise, when we measured nucleoid area, we saw an overall increase, indicating nucleoid decompaction in ΔpolyP cells during nitrogen starvation. It is therefore possible that the appearance of “high mobility” populations in intermediate and smaller particle sizes reflects an increase in the effective mesh size of the nucleoid, causing some particles to be less confined. Instead of being limited to the nucleoid-free regions of the cytoplasm, such small and intermediate particles may explore the nucleoid region more freely. These findings suggest that polyP modulates nucleoid compaction, and also raise the possibility that nucleoid compaction by polyP may be functionally important in globally affecting mobility of protein complexes during nitrogen starvation in a size-dependent manner.

### Limitations of the nucleoid area measurements

We use nucleoid origins to count the number of chromosomes per cell and we find a similar distribution of origin numbers in growing and starved ΔpolyP cells. However, it is possible that in the ΔpolyP strain a larger fraction of the cells have completely copied their chromosomes but not divided compared to exponentially growing cells. This would mean that starved ΔpolyP cells have on average more DNA per cell than growing cells, and this additional DNA could explain their larger nucleoid area. Yet in this case, we would then expect smaller cells, for which a large fraction (at least half) contain only one origin/chromosome per cell, to have similar nucleoid areas in ΔpolyP and WT cells. Instead, the trend we observe of larger nucleoid area in ΔpolyP cells is similar across all cell sizes, consistent with our interpretation that nucleoid decompaction is responsible for the larger nucleoid areas we measure.

### Distinct role of polyP in nitrogen but not carbon starvation

While polyP has a significant effect on cytoplasmic mobility during nitrogen starvation, we do not observe a similar effect during carbon starvation, suggesting distinct and context-dependent roles for polyP. Carbon and nitrogen starvation are both physiologically important states that lead to growth arrest, but they are also fundamentally different. Carbon starvation can lead to energy limitation in the absence of an inorganic electron donor for respiration. In contrast, nitrogen-starved cells, while lacking precursors for anabolism, are not intrinsically energy limited - although it is worth noting that regulatory coupling mechanisms whereby nitrogen limitation can inhibit carbon uptake may render nitrogen-starved cells effectively energy limited (29, 30). Nevertheless, we find that the effects of nitrogen and carbon starvation on cytoplasmic motion are qualitatively and quantitatively distinct in WT cells: under carbon starvation, we observe the emergence of a subpopulation of particles with decreased *D_app_* and *α*, which may be due to energy limitation. Decreased cytoplasmic motion under energy limiting conditions has been previously observed in both *Caulobacter crescentus* and *Escherichia coli* (11). Curiously, we also see the emergence of a second population with higher mobility than during exponential growth. Given that most cells have a single GFP-μNS particle, these particles may represent a sub-population of cells that are more metabolically active during this relatively early point in starvation. In contrast to carbon starvation, under nitrogen starvation while we also see partitioning of GFP-μNS particles into ‘low’ and ‘high’ mobility populations, the low mobility particles are of similar mobility in both unstarved and starved cells, meaning that overall the mobility of these particles has not decreased in starvation. In the ΔpolyP mutant, however, more particles have higher mobility, suggesting that polyP normally functions during early nitrogen starvation to suppress mobility to levels similar to what is observed in exponential growth. It seems therefore that polyP, made under both nitrogen and carbon starvation, may have distinct functions in these different states, with a specialized role during nitrogen starvation to modulate cytoplasmic mobility that is not necessary under the energy-limited carbon starvation state. Such specialization in nitrogen and carbon starvation has precedence: the other ancient master regulator of bacterial starvation responses, guanosine tetraphosphate (p)ppGpp, has been shown to have distinct roles in carbon and nitrogen starvation. Under nitrogen starvation, (p)ppGpp is required to slow transcription in order to ensure that transcription and translation are both attenuated in a coordinated fashion. Under carbon starvation, however, (p)ppGpp is not needed to attenuate transcription because it is already limited (31).

### Functional significance of tuning cytoplasmic mobility during growth-arrest

Under environmental conditions where one macronutrient is limited, but energy is not limited, such as nitrogen starvation, cells must still adjust many metabolic and regulatory processes in a concerted manner to arrest growth. PolyP biosynthesis occurs in response to diverse stress and starvation cues, but its molecular mechanism of promoting fitness in these different states has remained unclear. Polyphosphate has been shown to function as a non-specific protein chaperone, which could affect diverse cellular processes (26, 32). Our results suggest that polyP also plays a role in maintaining cytoplasmic mobility during nitrogen starvation. In the absence of polyP, a 2-fold effect on *D_app_* could modulate diffusion-controlled reactions *in vivo*, as could increased accessibility to the nucleoid of small protein complexes. How polyP contributes to nucleoid compaction during starvation remains to be determined. One possibility is that polyP granules, which form in the nucleoid region, may bring chromosomal regions together. A non-exclusive alternative is that soluble polyP can also drive nucleoid compaction. It is thought that in growing cells RNA may contribute to nucleoid compaction via crowding or charge-driven mixing of these polyanions ((23) and references therein). Given that transcription rates drop in response to growth arrest and polyP synthesis increases, one possibility is that polyP serves as a compensatory polyanion during starvation. Future work is needed to determine how polyP exerts these effects on the cytoplasm and nucleoid.

## Materials and Methods

### Strains

The SI Appendix contains tables S4, S5, S6 respectively of the strains, plasmids, and primers used in this study. The SI Appendix contains a detailed description of plasmid and strain construction.

### Cultivation conditions

The growth and starvation experiments were all performed in minimal media buffered with MOPS. From LB plates, strains were acclimated to the minimal media through overnight culture. 5mL subcultures were grown in glass test tubes into log phase at 37°C (OD500 = 0.4-0.6, takes 4.5 hours) and spun down, and the pellet was resuspended in MOPS-buffered media with either limiting nitrogen or limiting carbon for 6 h. For experiments with rhamnose-inducible GFP-μNS, 50μM rhamnose was added to the subcultures for the 4.5 hour growth period, but not added during the starvation period. Media composition is described in the SI appendix.

### Fluorescence Microscopy

Live cells were imaged at 26°C on 1% agarose pads containing MOPS-buffered minimal media for “0 h” data, and MOPS-buffered minimal media lacking added ammonium chloride for N-starvation experiments, or lacking sodium succinate for C-starvation experiments. Agarose pads were applied to 50 mm glass coverslip bottom dishes with plastic lids (Willco Wells, 0.17 mm/#1.5). Phase contrast and fluorescence images were acquired with a Nikon Ti2-E inverted microscope with perfect focus and the following other hardware: Objective: Plan apochromat phase contrast 100X oil immersion objective, N.A. 1.45, Illumination Source: For brightfield, a white LED, for fluorescence, the Spectra X Light Engine with a 470 nm LED (Lumencor) and a filter cube with a 466/40 nm bandpass excitation filter, 525/50 nm bandpass emission filter, and 495 nm dichroic mirror (Semrock). Camera: Prime 95B sCMOS with 11 μm x 11 μm pixel area (Photometrics). Image acquisition was controlled using Nikon Elements. For the chromosomal origin tracking experiments, the “ND Acquisition” module was used and images in phase and GFP fluorescence were acquired at 5s intervals for 100 frames, with the following parameters: For GFP, 100% light intensity from the LED, 100ms exposure time; for phase contrast, 75% light intensity and 100ms exposure time. For GFP-μNS particle tracking, the “Illumination Sequence” module was used to acquire GFP fluorescence using 30ms exposure time and either no delay, or a total of 210ms delay (including the 30ms exposure time) at 100% light intensity. A single phase contrast image was acquired at the beginning and end of the time series for the GFP-μNS particle tracking. For nucleoid imaging of HU-mCherry labeled cells, the Spectra X Light Engine with a 555 nm LED (Lumencor) and a filter cube with a 562/40 nm bandpass excitation filter, 641/75 nm bandpass emission filter, and 593 nm dichroic mirror (Semrock), 100% light intensity from the LED, 100ms exposure time. For nucleoid imaging of Sybr Green-labeled cells, live cells were labeled for 10 minutes with 1X Sybr Green and imaged immediately using the same configuration with GFP, but with 50 ms exposure time and the LED at 100% light intensity.

### Quantification of particle motion

We extracted particle trajectories and *MSD* curves from the raw movies using trackpy, an open-source tracking software written in python (17). Briefly, we used this algorithm to identify intensity maxima in each frame of a movie and to link them into particle trajectories throughout the movie. For each type of fluorescent spot (chromosome origins or GFP-μNS) we first tuned the input parameters to trackpy using a Jupyter notebook written for this purpose (18). We then applied the same parameters for all movies of the same type; for a table of our final input parameters to trackpy see SI Methods.

We extracted the values for the apparent diffusion coefficient, *D_app_*, and scaling exponent, *α*, from linear fits to the log-log curves of the mean–squared–displacement. For chromosome origins we applied the fit to the time interval 10–40 s and for μNS particles to the time interval 0.21–0.84 s, chosen such that the log-log curves are above the measured noise floor but still within their linear regime and away from noise due to the decreasing number of measurements at higher time lags. In this description *D_app_* is related to the y-offset of the fitted lines and *α* is their slope.

To estimate the noise level for our measurement of the *MSD*, we tracked movies of fixed cells as described above, then used the resulting ensemble mean-squared-displacement (*eM SD*) as the noise floor shown in Figures 1, 2. We calculated this *eM SD* using trackpy, after the expression

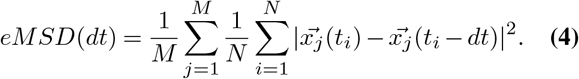

where *M* is the total number of spots, *N* is the number of statistically independent measurements for a given locus and *dt*, and 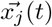 is the position of locus j at time *t* (see (17) for more details). We find that the *eM SD* varies slightly among different strains and conditions; for the 0 h data and the 6 h data in nitrogen starvation we show the *eM SD* obtained from fixed ΔpolyP cells 6 h into starvation as the most conservative limit among these four conditions.

### Size estimation for GFP-μNS particles

We estimated the size of GFP-μNS particles from the integrated intensity of their fluorescent spots, following an approach similar to (11). First, we applied a Gaussian fit to the intensity profile of all bright spots using the expression

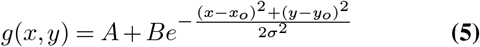

where *x_o_, y_o_* are the particle’s coordinates, *A* is the local intensity background, *B* the particle’s intensity amplitude and *σ* is its width; see SI Methods for a list of the initial guess for each parameter. We omitted very large particles with width 2*σ*^2^ > 450 nm (~ 2 × our diffraction limit) as they are likely either out of focus or subject to motion blur. As an indicative value for the width we took the average *σ* over the first five frames of the particle’s appearance. From the results of these fits we calculated the total intensity of each particle as the integral of its Gaussian intensity profile, *I* = 2*πBσ*^2^, and we estimated its size, *d*, as 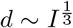, where for *I* we took, as before, the average value from the first five frames of the particle’s appearance.

In all tracking analysis we omitted particles for which any one of the parameters we calculated is non-physical, i.e. nonpositive, or whose *MSD* cannot be fitted.

### Estimation of the nucleoid-cytoplasm (NC) ratio

We obtained the outlines of cells from phase images using MicrobeTracker or Oufti (24, 33). For the datasets where we used Microbetracker we imported the results for cell segmentation into Oufti (24). We segmented the nucleoid, labeled with HU-mCherry, using the Object Detection module; for a list of input parameters see SI methods.

Except for segmentation we performed all analysis in python. Our scripts as well as step-by-step instructions to apply our analysis pipeline are available in (18).

## ACKNOWLEDGEMENTS

We thank the Center for Environmental Microbial Interactions (CEMI) at Caltech, where the project was conceived. We are grateful to all members of the Manley and Racki groups for insightful discussions, and to Ravi Chawla for testing our tracking parameters on carbon starvation data. Funding was received from the European Research Council (ERC CoG 819823, Piko to Su.M. and So.M.), the Swiss National Science Foundation (182429 to Su.M. and W.S.), and the Donald E. and Delia B. Baxter Foundation Fellowship (L.R.R.) This is manuscript #30145 from The Scripps Research Institute.

## Supplementary Information

### Quantification of particle motion

We localized and tracked chromosome origins using trackpy 0.4.2 (17), an open-source software package based on the particle-tracking algorithm originally described in (34). From the resulting trajectories we calculated mean-squared-displacement curves and used a least-squares fit to extract values for the diffusion coefficient and for the scaling exponent. We performed all data analysis in python. We fit using the ‘optimize’ package of scipy, and we tested the statistical significance of differences in the means or medians of the resulting distributions using the ‘stats’ package of scipy.

To facilitate the process of identifying suitable tracking parameters, we have integrated trackpy and complementary functions in a Jupyter Notebook (18).

Typical values we used are shown in Table S1. The diameter was obtained by measuring the typical width of a bright spot in Fiji and ensuring there is no sub-pixel bias in tracking (see also trackpy tutorial (17)); the minmass was chosen by trial-and-error on a few sample frames; the search range was chosen as the width of a cell, after verifying that it is beyond the range of the step size distribution of our trajectories; memory and stub length were chosen by trial-and-error, looking at sample tracks. We excluded all spots with SNR < 1.

We noticed that on dense fields of view the height of the *D_app_* distributions increased at higher values. We hypothesize that this is due to differences in the pre-processing of the image: in denser regions the estimated local background may be higher due to larger amounts of cytosolic background. When this higher background is subtracted, smaller particles can be better discerned and might contribute to additional counts at high *D_app_*. To mitigate this effect, we omitted from the analysis movies where bacteria were closely packed in the field of view.

**Table S1.**
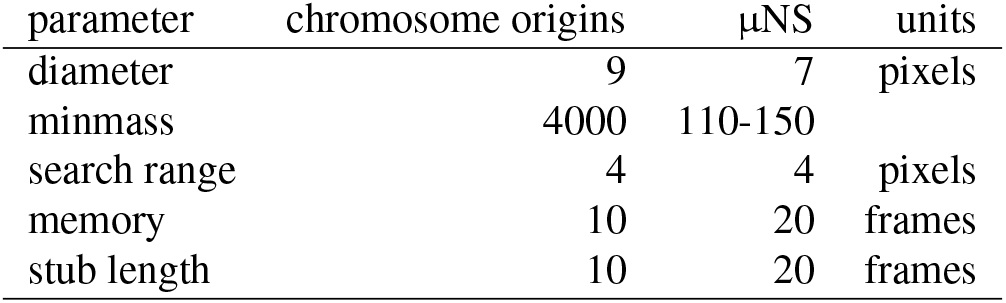
Input parameters to trackpy for particle tracking.

**Table S2.** Initial guess for Gaussian fits to calculate the integrated intensity of μNS particles.

| parameter | initial guess | units |
| --- | --- | --- |
| offset | 100 |  |
| amplitude | 100 |  |
| $x_o, y_o$ | coordinates obtained from trackpy | pixels |
| $\sigma$ | 3 | pixels |

### Estimation of the nucleoid-cytoplasm ratio

We segmented the cells in Microbetracker and the nucleoid in Oufti using the ObjectDetection feature (24). For a list of input parameters in ObjectDetection, see Table S3. We selected most parameters by trial and error, following the values quoted in (22). We removed cases where the nucleoids of two cells were mistakenly segmented as one continuous nucleoid, as these were cases where the cells were touching due to how they had landed on the pad and should not lead to a biased selection. The sigma of PSF in ObjectDetection corresponds to our imaging system (NA = 1.45, *λ* = 600 nm, 1 pixel = 0.10748 *μm*).

### Plasmids

The new plasmids used in this study were generated using restriction enzyme digestion and ligation or Gibson Assembly (35). Plasmid pJM220 is a derivative of pUC18R6K-mini-Tn7T-Gm and was generated as described previously (36), by amplifying rhaSR-P_rhaBAD_ from *E. coli* (We used *E. coli* strain MG1655 as a template, which has the identical sequence of these genes as the strain used in the original study, W3110) using primers LRPR873F and LRPR874R. Both the plasmid backbone and PCR product were cut with PstI and SacI and ligated. Plasmid pLREX119 is a derivative of plasmid pJM220, and was generated by inserting the coding sequence of GFP-μNS (See SD-GFP-μNS Sequence, as designed by (11), but with the addition of a Shine Dalgarno sequence). The GFP-μNS sequence was ordered from IDT as a gBlock and amplified with linkers by the primers LRPR879F, LRPR884R. This sequence was inserted into plasmid pJM220, cut with restriction enzymes SpeI and PstI. All plasmids were validated by sequencing.

**Table S3.** Input parameters for nucleoid object detection in Oufti.

| parameter | value |
| --- | --- |
| manual background threshold | 0.2 |
| background subtraction method | 3 |
| background subtraction threshold | 0.1 |
| background filter size | 8 |
| smoothing range (pixels) | 2 |
| magnitude of LOG filter | 0.1 |
| sigma of PSF | 0.82 |
| fraction of object in cell | 0.4 |
| minimum object area | 40 |

### Strains

The unmarked deletion strain referred to as ΔpolyP in this study was constructed previously (16), and is a quadruple knockout of *P. aeruginosa*’s four polyP kinases, Ppk1 (PA14_69230), Ppk2A (PA14_01730), Ppk2B (PA14_33240), and Ppk2C (PA14_19410). Strains LR231 and LR238 were used both for chromosomal origin counting and tracking, and were constructed previously (37). Reporter strains with insertions at the *att*Tn7 site were generated by tetraparental conjugation, and exconjugants were selected on VBMM medium with 100mg/mL gentamycin, as described previously (38), and verified by PCR.

### Media and growth conditions

Strains were grown at 37°C shaking at 250 rpm in “Complete” MOPS-buffered minimal media (40mM sodium succinate, 22mM NH_4_Cl, 43mM NaCl, 2.2mM KCl, 1.25mM NaH_2_P0_4_, 1mM MgSO_4_, 0.1mM CaCl_2_, 7.5μM FeCl_2_•4H_2_O, 0.8μM CoCl_2_•6H_2_O, 0.5μM MnCl_2_•4H_2_O, 0.5μM ZnCl_2_, 0.2μM Na_2_MoO_4_•2H_2_O, 0.1μM NiCl_2_•6H_2_O, 0.1μM H_3_BO_3_, and 0.01μM CuCl_2_•2H_2_O, 50mM MOPS, pH 7.2). 5mL cultures at OD500 = 0.0125 to 0.025 were grown in glass test tubes to OD500 = 0.4 to 0.6, then spun down at room temperature in at 5000xG, and resuspended to OD500 = 0.4 in nitrogen-limited or carbon-limited MOPS-buffered minimal media (Identical to MOPS-buffered media, but with 1mM NH_4_Cl instead of 22mM for nitrogen-limited media, and with 1mM sodium succinate instead of 40mM sodium succinate for carbon-limited media respectively) in clean test tubes. For exponentially growing cells, i.e. “0h,” cells were either collected and imaged before being spun down and shifted to nitrogen-limited medium, or they were back-diluted in Complete MOPS-buffered minimal media to OD500 = 0.025 and allowed to continue growing for 2h before imaging.

### SD_GFP-μNS Sequence

AGGAGGATATACATATGGTGAGCAAGGGCGAGGAGCTGTTCACCGGGGTGGTGCCCATCCTGGTCGAGCTGG ACGGCGACGTAAACGGCCACAAGTTCAGCGTGTCCGGCGAGGGCGAGGGCGATGCCACCTACGGCAAGCTGA CCCTGAAGTTCATCTGCACCACCGGCAAGCTGCCCGTGCCCTGGCCCACCCTCGTGACCACCCTGACCTACGG CGTGCAGTGCTTCAGCCGCTACCCCGACCACATGAAGCAGCACGACTTCTTCAAGTCCGCCATGCCCGAAGGC TACGTCCAGGAGCGCACCATCTTCTTCAAGGACGACGGCAACTACAAGACCCGCGCCGAGGTGAAGTTCGAGG GCGACACCCTGGTGAACCGCATCGAGCTGAAGGGCATCGACTTCAAGGAGGACGGCAACATCCTGGGGCACA AGCTGGAGTACAACTACAACAGCCACAACGTCTATATCATGGCCGACAAGCAGAAGAACGGCATCAAGGTGAA CTTCAAGATCCGCCACAACATCGAGGACGGTTCTGTTCAGCTGGCGGACCACTACCAGCAGAACACCCCGATC GGTGACGGTCCGGTTCTGCTGCCGGACAACCACTACCTGAGCACCCAGTCCGCCCTGAGCAAAGACCCCAACG AGAAGCGCGATCACATGGTCCTGCTGGAGTTCGTGACCGCCGCCGGGATCACTCTCGGCATGGACGAGCTGTA CAAGTCCGGACTCAGATCTCGAGCTCAAGCTTCGAATTCTGCAGTCGACTCTAGAGGATCCGTCATGGCTTCC AATGACGTGACAGATGGGATCAAGCTTCAGTTGGACGCATCTAGACAGTGTCATGAATGTCCTGTGTTGCAGC AGAAAGTGGTTGAGTTAGAGAAACAGATTATTATGCAGAAGTCAATCCAGTCAGACCCTACCCCAGTGGCGCTG CAACCATTGTTGTCTCAGTTGCGTGAGTTGTCTAGTGAAGTTACTAGGCTACAGATGGAGTTGAGTCGAGCTCA GTCCCTGAATGCTCAGTTGGAGGCGGATGTCAAGTCAGCTCAATCATGTAGCTTGGATATGTATCTGAGACACCA CACTTGCATTAATGGTCATGCTAAAGAAGATGAATTGCTTGACGCTGTGCGTGTCGCGCCGGATGTGAGGAGAG AAATCATGGAAAAGAGGAGTGAAGTGAGACAAGGTTGGTGCGAACGTATTTCTAAGGAAGCAGCTGCCAAATG TCAAACTGTTATTGATGACCTGACTTTGATGAATGGAAAGCAAGCACAAGAGATAACAGAATTACGTGATTCG GCTGAAAAATATGAGAAACAGATTGCAGAGCTGGTGAGTACCATCACCCAAAACCAGATAACGTATCAGCAAGA GCTACAAGCCTTGGTAGCGAAAAATGTGGAATTGGACGCGTTGAATCAGCGTCAGGCTAAGTCTTTGCGTATTA CTCCCTCTCTTCTATCAGCCACTCCTATCGATTCAGTTGATGATGTTGCTGACTTAATTGATTTCTCTGTTCCAACT GATGAGTTGTAA

**Table S4.** Strains

| name | genotype | description | source |
| --- | --- | --- | --- |
| *P. aeruginosa* strains |  |  |  |
| LR31; DKN263 | *P. aeruginosa* UCBPP-PA14 | wild type background for all other strains | Ref. (16) |
| LR119; DKN1731 | *P. aeruginosa* UCBPP-PA14 $\Delta ppk1$ $\Delta ppk2A$ $\Delta ppk2B$ $\Delta ppk2C$ | deletion of PA14_69230, PA14_01730, PA14_33240, and PA14_19410 in DKN263 | Ref. (16) |
| LR231 | *P. aeruginosa* UCBPP-PA14 $attTn7::mini-Tn7T-Gm^R$ $ParS^{pMT1}$ $P_{ssb}::ssb-mCherry$ $gfp-paBr^{pMT1}$ | Wt UCBPP-PA14 with $ParS^{pMT1}$ motif ($parB^{pMT1}$ protein binding site) and both $ssb-mCherry$ and $gfp-ParB^{pMT1}$ chimeras expressed under control of the $ssb$ promoter, all integrated at the $attTn7$ site on the chromosome | Ref. (37) |
| LR238 | *P. aeruginosa* UCBPP-PA14 $\Delta ppk1$ $\Delta ppk2A$ $\Delta ppk2B$ $\Delta ppk2C$ $attTn7::mini-Tn7T-Gm^R$ $ParS^{pMT1}$ $P_{ssb}::ssb-mCherry$ $gfp-parB^{pMT1}$ | $\Delta polyP$ UCBPP-PA14 with $ParS^{pMT1}$ motif ($parB^{pMT1}$ protein binding site) and both $ssb-mCherry$ and $gfp-ParB^{pMT1}$ chimeras expressed under control of the $ssb$ promoter, all integrated at the $attTn7$ site on the chromosome | Ref. (37) |
| LR453 | *P. aeruginosa* UCBPP-PA14 $attTn7::mini-Tn7T-Gm^R$ $rhaSR-P_{rhaBAD}::gfp-\mu NS$ | Wt UCBPP-PA14 with rhamnose-inducible GFP-$\mu NS$ integrated at the $attTn7$ site on the chromosome; made using pLREX119 | This study |
| LR454 | *P. aeruginosa* UCBPP-PA14 $\Delta ppk1$ $\Delta ppk2A$ $\Delta ppk2B$ $\Delta ppk2C$ $attTn7::mini-Tn7T-Gm^R$ $rhaSR-P_{rhaBAD}::gfp-\mu NS$ | $\Delta polyP$ UCBPP-PA14 with rhamnose-inducible GFP-$\mu NS$ integrated at the $attTn7$ site on the chromosome; made using pLREX119 | This study |
| *E. coli* strains |  |  |  |
| DKN1298 | *E. coli* SM10$\lambda pir$, pTNS1 | Helper strain for integration at the $attTn7$ site | Ref. (38) |
| DKN1299 | *E. coli* HB101, pRK2013 | Helper strain for conjugation | Ref. (38) |
| LR448 | *E. coli* TurboCells, pJM220 |  | Ref. (36) |
| LR456 | *E. coli* TurboCells, pLREX119 |  | This study |

**Table S5.** Plasmids

| name | description | source |
| --- | --- | --- |
| pJM220 | *pUC18T-miniTn7T-Gm<sup>R</sup>-rhaSR-P<sub>rhaBAD</sub>*; chromosomal integration (*attTn7* site) and expression (rhamnose-inducible) vector | Ref. (36) |
| pLREX119 | *pUC18T-miniTn7T-Gm<sup>R</sup>-rhaSR-P<sub>rhaBAD</sub>:gfp-μNS*; for integration at *attTn7* site of rhamnose-inducible GFP-μNS chimera | This study |

**Table S6.**
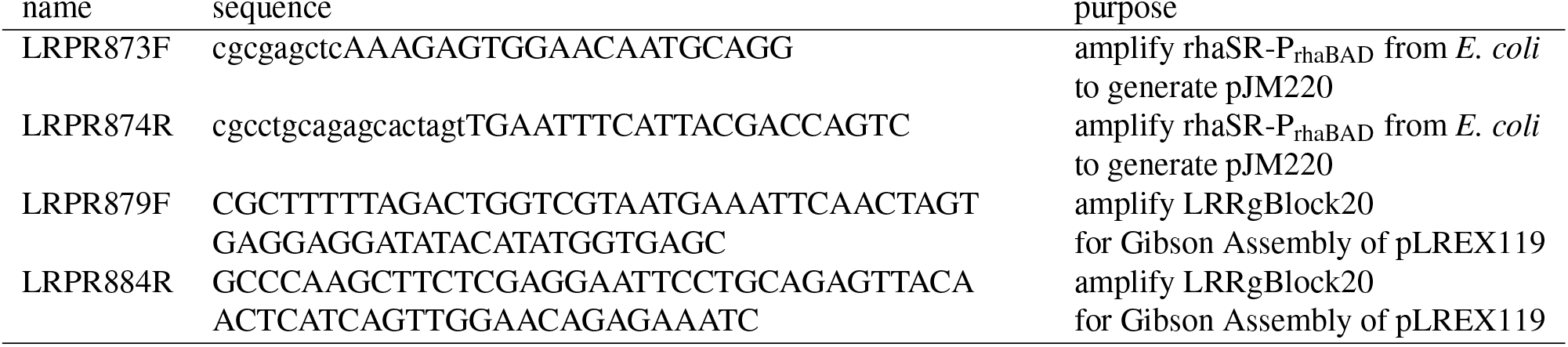
Primers

**Fig. S1.**
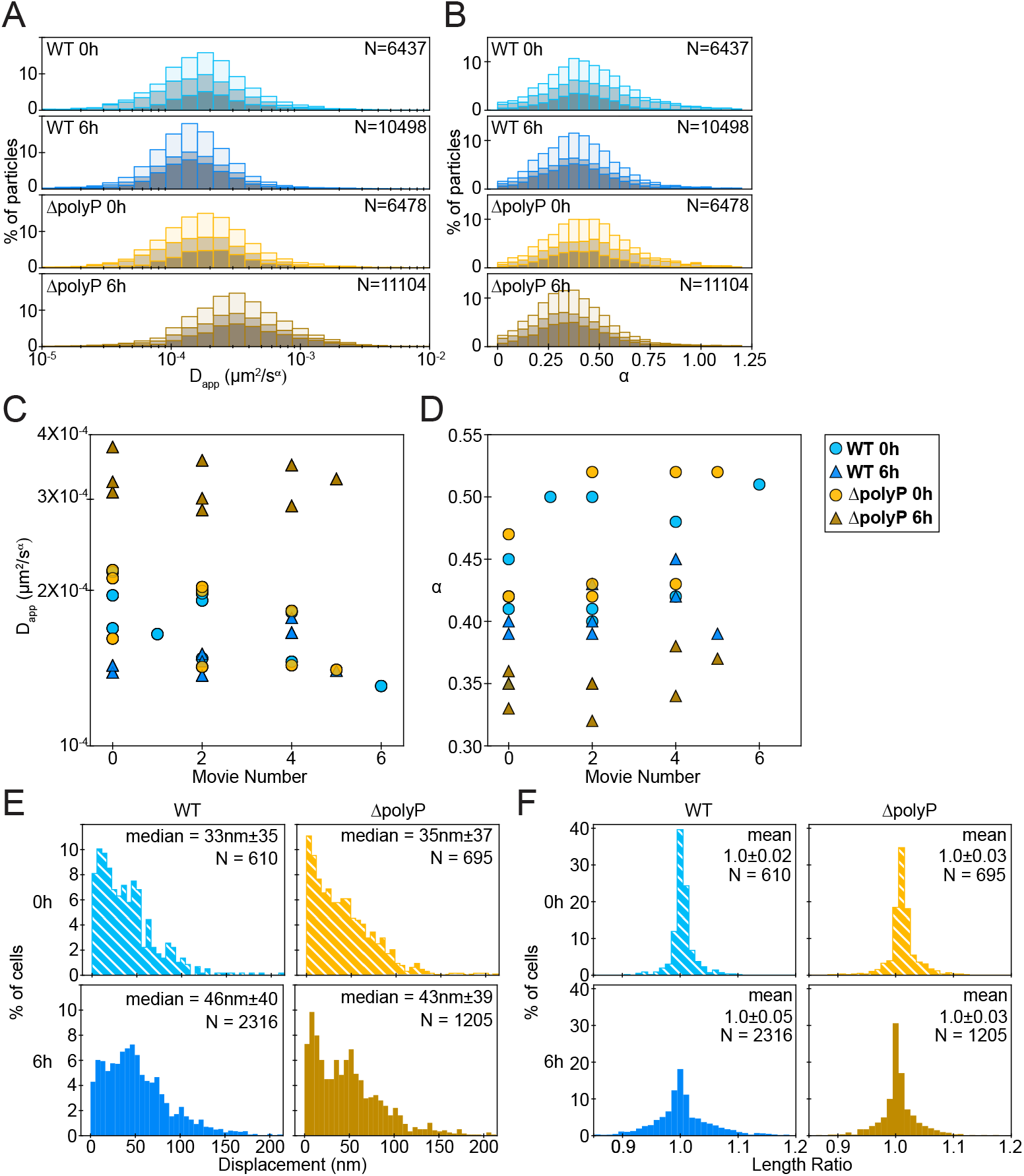
The mobility of chromosome origins 6 h into nitrogen starvation is not affected by experimental day or time on agarose pad. (A) Stacked histograms of *D_app_* for chromosomal origins in WT (blue) and ΔpolyP (orange) cells, 0 h and 6 h into N-starvation. For each condition, each sub-histogram, colored by shades of tinted gray, corresponds to data from a single day; we repeated this experiment on three different days. (B) Same as in (A) for *α*. (C) Median *D_app_* vs movie number for all recordings; we recorded several movies on each of the three days. Here the movie number corresponds approximately to the time since cells were plated on agarose pads for imaging. (D) Same as in (C) for *α*. (E) Histograms of the displacement of the cells’ center of mass between the first and the last frame. (F) Histograms of cell growth during the movies, measured as the ratio of each cell’s length at the last frame divided by its length at the first frame.

**Fig. S2.**
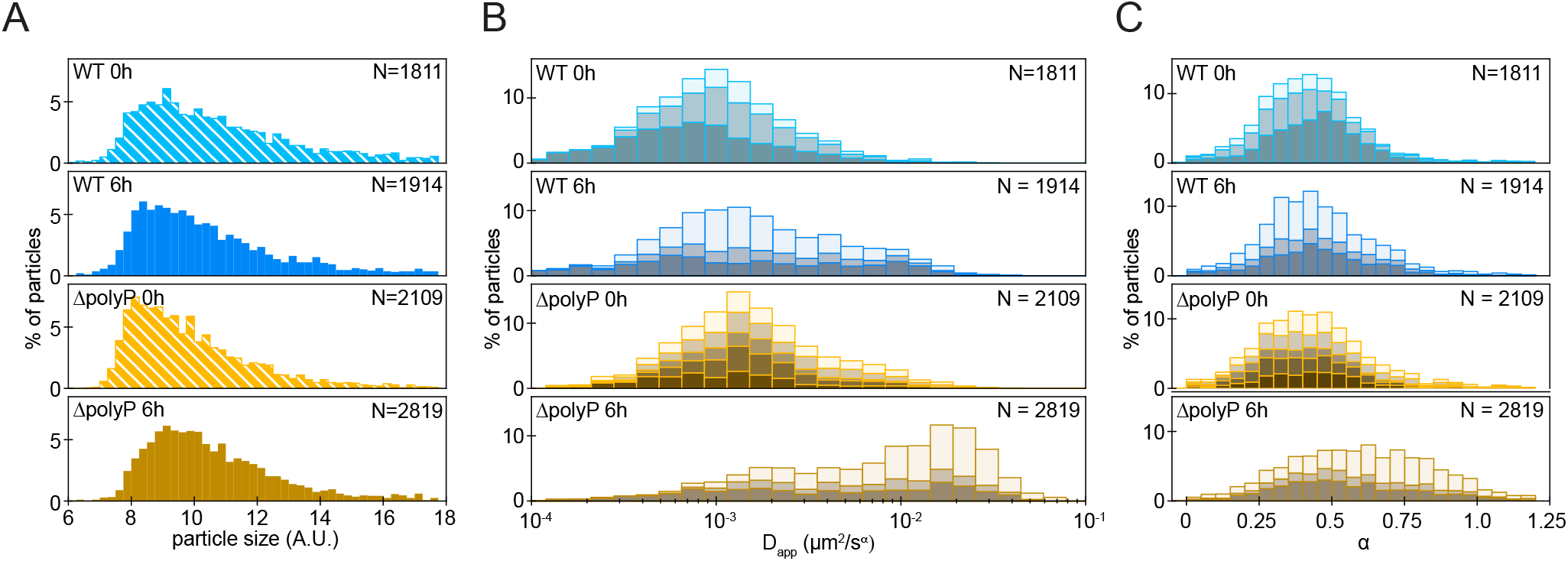
GFP-μNS particle size distribution and day-to-day variability in mobility. (A) Particle size distribution for GFP-μNS. (B) Stacked histograms of *D_app_* for GFP-μNS particles in WT (blue) and ΔpolyP (orange) cells, 0 h and 6 h into N-starvation. For each condition, each sub-histogram, colored by shades of tinted gray, corresponds to data from a single day. We repeated this experiment on five different days in total; because we excluded from the analysis movies with a dense field of view, as explained in the SI methods (Quantification of particle motion), for most conditions we have included data from three of those days. (C) Same as in (B) for *α*.

**Fig. S3.**
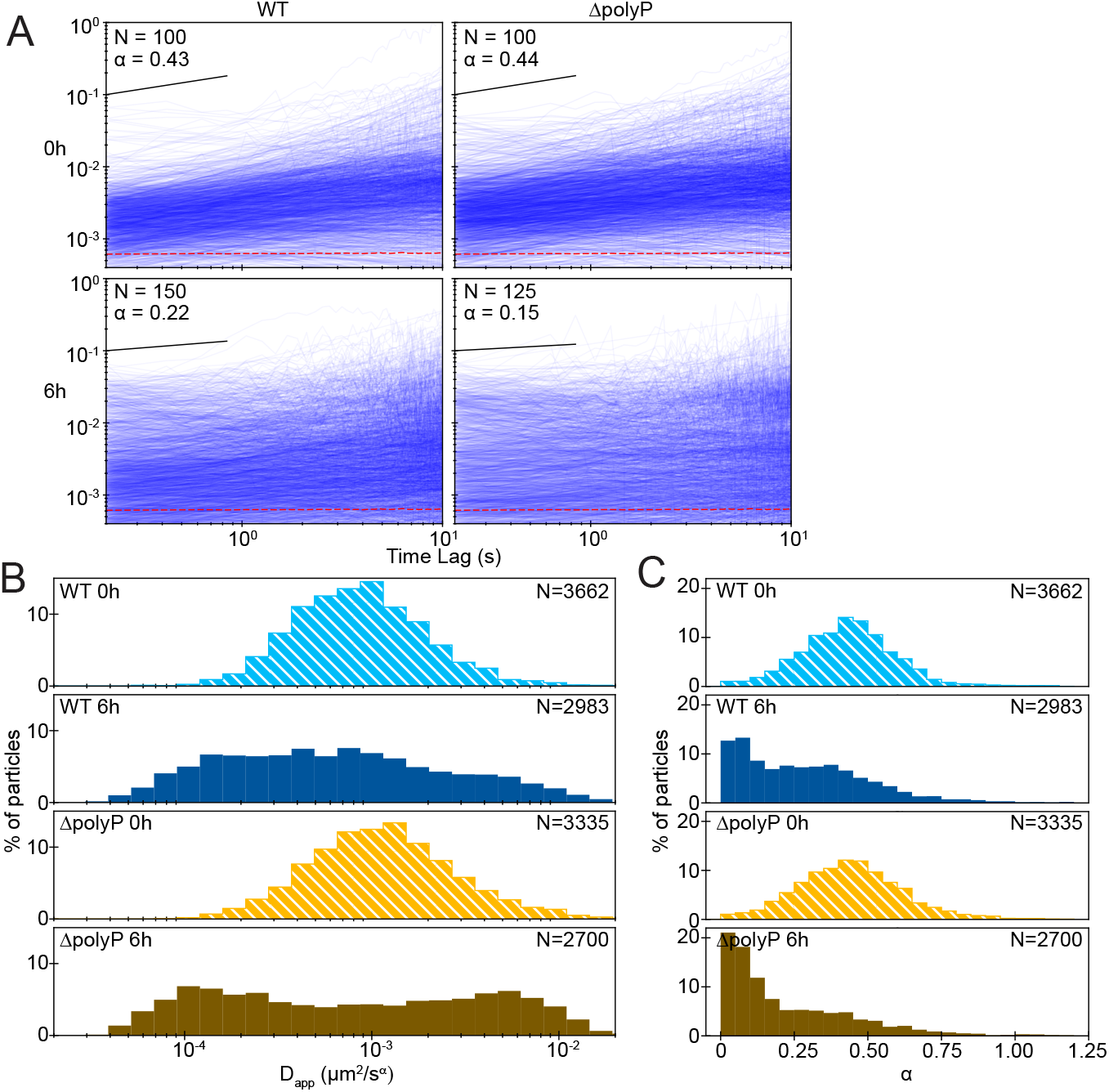
Effect of polyphosphate on the mobility of μNS particles under carbon starvation. (A) Examples of mean-squared-displacement curves for a randomly chosen subset of particles, 0 h (top row) and 6 h into C starvation (bottom row), for WT (left column) and ΔpolyP cells (right column). The *α* values in each panel denote the median of the distribution; they were obtained from linear fits over the time interval spanned by the black line (0.21 s – 0.84 s). N is the number of plotted curves; we show a randomly chosen subset for clarity. The red dotted line indicates our empirical noise floor, measured as the ensemble mean-squared-displacement of GFP-μNS particles in fixed cells. (B) Histograms of *D_app_* values extracted from the *MSDs* shown in (A). (C) Histograms of *α* values extracted from the *MSDs* shown in (A). In both (B), (C), N indicates the total number of spots.

**Fig. S4.**
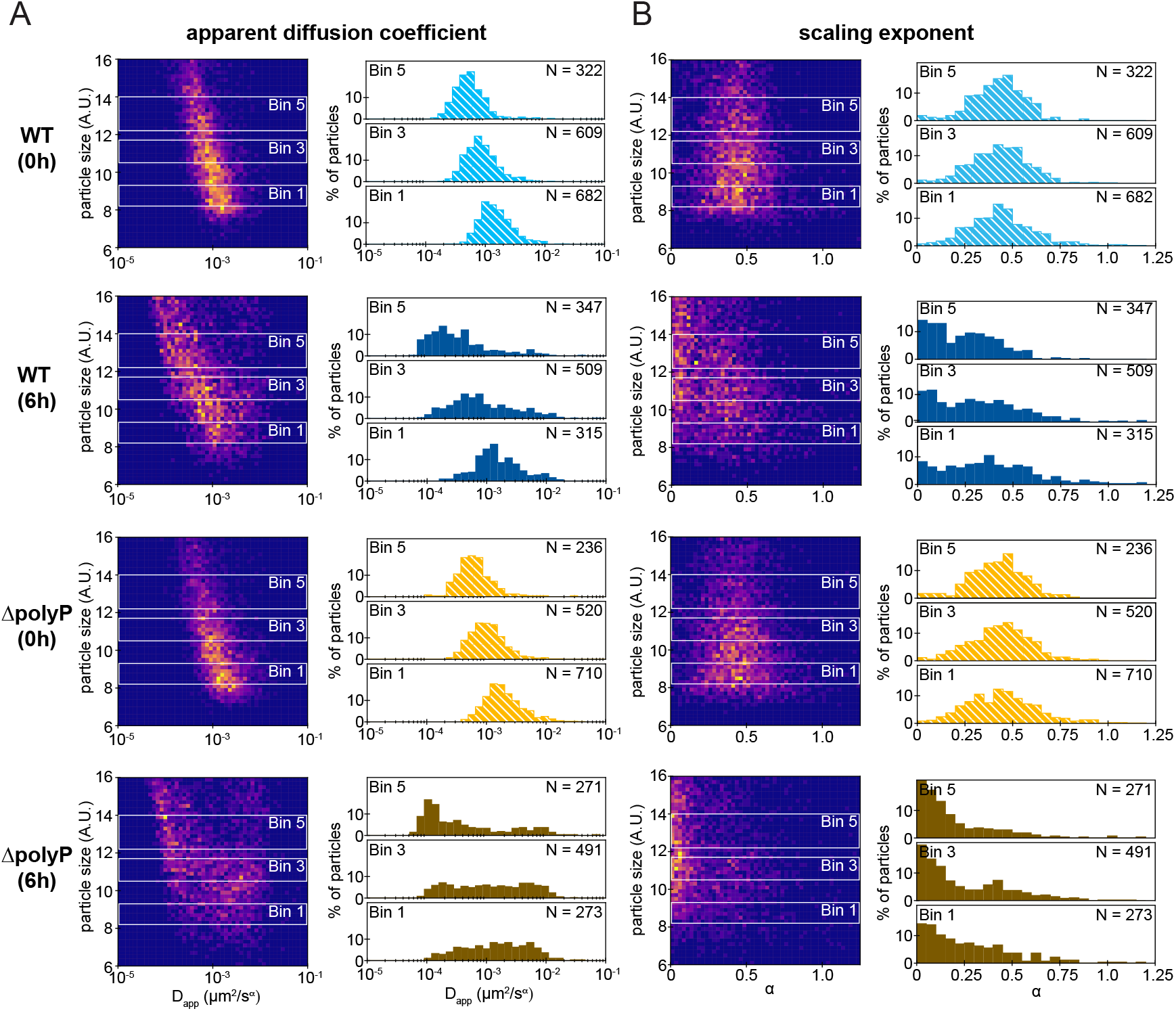
GFP-μNS particle size and cytoplasmic mobility under carbon starvation. (A) Left: heatmap of particle size vs *D_app_*; right: histograms of the *D_app_* distribution for three size bins in each condition. (B) Left: Heatmap of particle size vs *α*; right: histograms of the *α* distribution for three size bins in each condition. Bin 1 corresponds to small particles, bin 3 to medium-sized particles, and bin 5 to large particles.

**Fig. S5.**
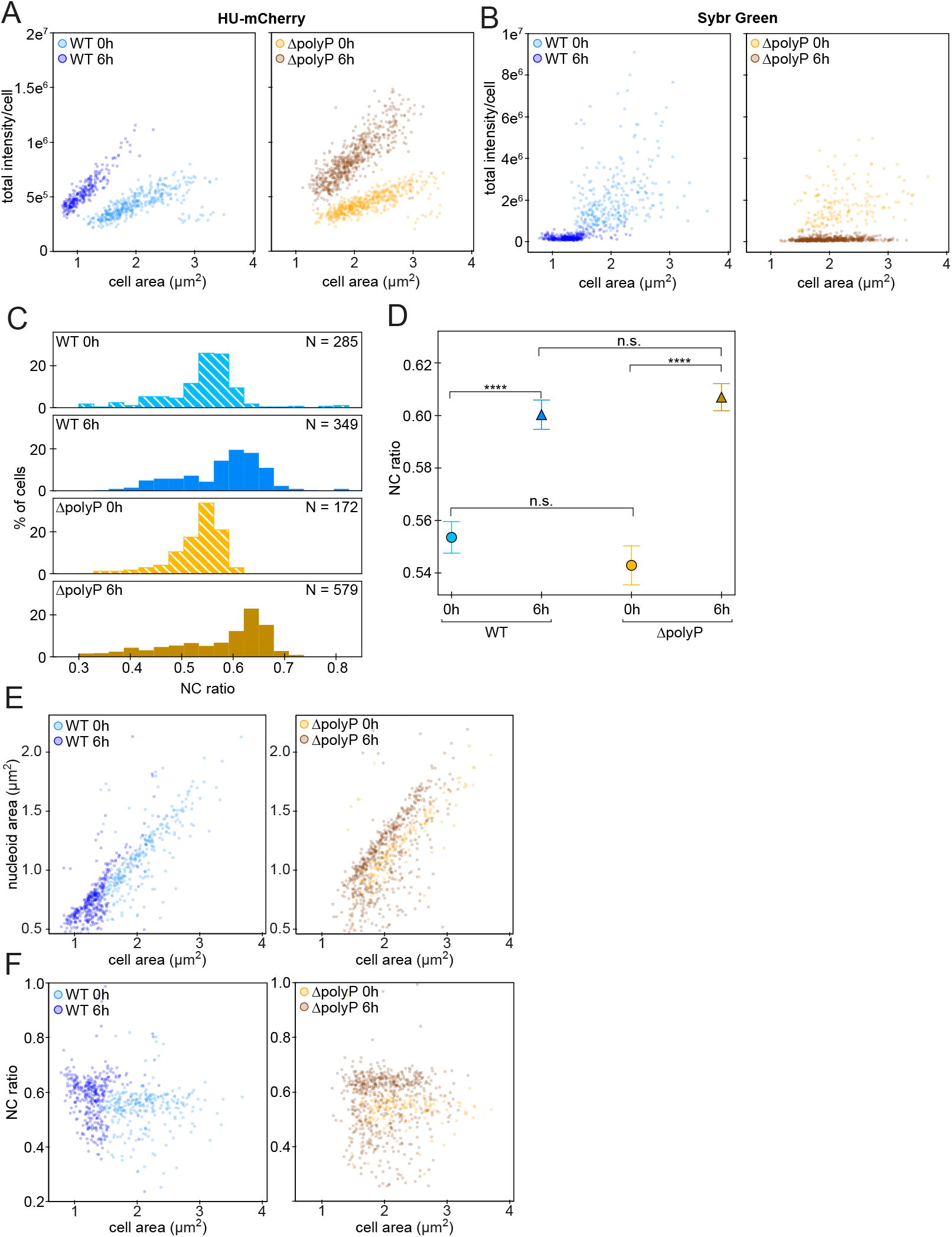
Controls for nucleoid area using Sybr Green. (A) Total fluorescence as a function of cell area for HU-mCherry-labeled cells. (B) Total fluorescence as a function of cell area for Sybr Green-labeled cells. (C) Histograms of the NC ratio for all four conditions for Sybr Green-labeled cells (D) Medians of the distributions of the NC ratios shown in (C). The error bars denote the standard error of the mean, and the asterisks denote the statistical significance of the differences in the medians, according to the Mood’s median test. (E) Scatter plots of nucleoid area vs cell area for Sybr Green-labeled cells. (F) Scatter plots of NC ratio vs cell area for Sybr Green-labeled cells.

